# Video-Language Models as Flexible Social and Physical Reasoners

**DOI:** 10.1101/2024.05.11.593700

**Authors:** Daniel Fleury

**Affiliations:** Johns Hopkins University

## Abstract

From an early age, humans are challenged with evaluating rich environments full of socially and physically grounded concepts. For example, we might be spectating a rapidly unfolding tennis match, anticipating ball trajectories based on players’ body cues and goals. In another scenario, we may engage with long storylines, juggling the mental states of characters with varying knowledge of an unfolding conflict. The complexity of this learning problem is notable as it can be multimodal, integrate information at varying timescales, and implicitly co-attend to social and physical scene properties for downstream reasoning. Large language-vision models like GPT4-V, LLaMA-3, which use vision-language embeddings, show skills in commonsense psychology and physics, though they only process single images. Models like CLIP and VisualBERT encode visual information in high-level cortical areas but do not inherently capture video-level representations. This paper introduces a novel video-language architecture that incorporates pooled video embeddings into LLMs by first extracting spatiotemporal embeddings and mapping them to the model’s decoder through a learnable linear layer. We enhance the model by training it with video-caption pairs from the ADEPT and AGENT datasets, aimed at quantifying “surprisal” in physical and psychological contexts. Finally, we design separate voxel wise encoding models for videos involving physics and psychology using the hidden states and logits from the LLMs last layer and pre-projected CLIP embeddings. We find that hidden state activations can remarkably explain high variance (R^2 up to ∼70%) across dorsal physics regions and highly distributed, ventral social vision areas. Notably, for models trained to only encode physically surprising stimuli, the hidden states and *pre*-projected CLIP embeddings explain nearly identical regions of variance across the inferior-parietal lobule. However, when the encoding model is trained to encode only socially surprising events, hidden states explain far more distributed ventral and dorsal activations over pre-projected CLIP embeddings.

## Introduction

Navigating and reasoning about a highly visual world requires grounding our perception in basic intuitions of how agents, objects, and scenes are expected to interact across time. Although computationally separable on the surface, it becomes apparent that our psychological intuitions of agents and their interactions can saliently interact with our internal “physics engine.” Indeed, in infants as young as ten months of age, abstract variables such as “force,” “work”, or “effort” can be used to model their inference about agents’ goal preferences in videos with simple shapes (Liu et al., 2017). Relevantly, unified computational frameworks using deep reinforcement learning (RL) have been developed to render many Heider-Simmel-type stimuli in a simulated psychological space that captures a partition between physical law violations and differential involvement of social intention (Shu et al., 2021). In the same vein, converging behavioral and neurofunctional evidence suggest that the computations underlying both mental spaces are largely dissociable, regardless of how they are jointly represented in downstream tasks (Sajjad et al., 2023)(Maróthi & K′eri, 2014)(Mitko & Fischer, 2020)(Kamps et al., 2017). In adults with Williams syndrome (WS), a genetic developmental disorder that severely impairs spatial cognition, intuitive physical judgments were significantly worse than psychological judgments (Kamps et al., 2017). Further, event-related fMRI passive viewing paradigms find that social vs. physical categories with Heider-Simmel stimuli could be decoded in the pSTS/TPJ, with high correlation to the MT+ area/lateral occipital cortex (LOC) (Sajjad et al., 2023).

These results are heavily consistent with the localization of distinct theory of mind (ToM) and intuitive physics networks. The former primarily includes the bilateral temporal parietal junctions (TPJ), posterior superior temporal sulcus (pSTS), medial prefrontal cortex (mPFC), posterior cingulate cortex (PCC), and precuneus (Mason & Just, 2009) which are variably involved in representing simple to abstract social features along the lateral visual pathway (McMahon et al., 2023) (Isik et al., 2017). The latter encompasses the parieto-frontal network, chiefly centered on the supramarginal gyrus (SMG) which supports a wide range of computations involving manipulable object representations and action planning (Fischer & Mahon, 2022) (Fischer et al., 2016). Just as crucially, previous research has determined that climbing up the cortical hierarchy corresponds to integrating multimodal information across increasingly large temporal receptive windows (TRWs) (Lerner et al., 2011). For instance, brain responses in intermediate and high-level cortical regions (e.g. pSTS, TPJ, mPFC, and precuneus) only evoke reliable responses to spoken stories with intact sentence-level and paragraph-level sequences, respectively (Hsiang-Yun & Honey, 2020). Similar gradients of temporal integration also appear when viewing natural movies (Meer et al., 2020). Accordingly, computational modeling that attempts capturing humans’ (1) hierarchically sensitive (i.e. with respect to time and semantics) social perception capabilities along the ventral stream and (2) action-guided inference about future physical states in the dorsal stream are relevant.

Previously, physical violations of expectation (VoE) have been integrated into deep learning models by combining perception and dynamics modules (Piloto et al., 2022). Inspired by infants’ VoE when viewing physically unlikely scenarios, the model uses a perceptual module with unsupervised autoencoders to build up object representations of simple 3D shapes. Next, a dynamics predictor feeds latent object codes across time into a long-short term memory network (LSTM) in order to predict the next frame’s object code. Prediction error between predicted vs. actual object codes are then used as a proxy of “surprisal” when tested on physical concepts like continuity, object persistence, solidity, and inertia. Still, the extent to which feedforward vs. feedback strategies best model cortical processing remains at an impasse – and it is likely that a mixture of both are used. Top-down approaches may draw on Bayesian sampling to simulate likely physical outcomes (Battaglia et al., 2013) while bottom-up strategies employ sequence prediction of probable next event states based on object representations in memory (Li et al., 2024). Previously, psychological intuitions about how agents plan and act in accordance to goals and their world state have been simulated with Bayesian inverse planning and theory of mind neural networks that largely hinge on top-down connectivity (Shu et al., 2021). But the extent to which regions along social processing streams in the brain can be more accurately modeled by bottom-up discriminative models versus top-down inverse generative models is still an area of ongoing research (McMahon & Isik, 2023). Recent work suggests that bottom-up models can encode social interactions across the lateral visual stream while top-down models are more effective at mapping neural hubs for mentalization about agents’ beliefs, goals, and actions (Baker et al., 2011).

Importantly, the scope of this paper is *not* to discern the compatibility of feedforward versus generative models within intuitive physics and psychology networks. Instead, we note a gap in model-brain alignment in explaining the separability of these networks. That is, current research does not explore how multimodal video-language models can differentially encode social and physical expectations in a joint representational space. Further, current research largely overlooks how video embeddings encode brain activity before and after they are projected into the decoder space of modern LLMs. Models like contrastive language-image pretraining (CLIP) use natural language supervision in order to create an embedding space that maximizes the cosine similarity between joint image-text representations (Radford et al., 2021). Separate image and text encoders are jointly trained over many *n* pairs with an objective function of maximizing similarity across the diagonal of the dataset matrix. Moreover, this training process minimizes distance between pairs not in the diagonal. This is different from supervised CNNs in that the optimization goal is not to map image features to text labels in a feedforward manner, but rather predict correct embedding pairs. The result is a natural multimodal embedding space that represents gradients in vision-language alignment without exhausting as much computational resources (Joshi et al., 2024).

### Video-Language Models as Models of Commonsense Physics and Social Perception

Not surprisingly, efforts at assessing whether this embedding space can encode the cortical hierarchy have been explored in depth. Previous voxel wise encoding models using CLIP with a ResNet50 visual encoder accurately predict activity across the high-level visual cortex (e.g. PPA, EBA, and RSC) while outperforming the unimodal ResNet50 architecture alone (Wang et al., 2023). Further exploration shows that the V2 and V3 are better predicted by multimodal models, with higher layers in *image* transformers (i.e. unimodal) correlating with late visual areas and vice versa (Oota et al., 2022). It is also suggested that CLIP’s learning process may capture how abstract concepts penetrate early vision in a top-down manner. However, brain encoding models that use multimodal image-text transformers and CLIP embeddings alone are limited in two respects. First, both the fMRI datasets and multimodal embeddings in current studies only capture representations from single image-text pairs. Although datasets like BOLD5000 are naturalistic and categorically diverse, humans perceive interactions between objects, agents, and their environment with respect to time. Moreover, there is rich causal information within and between events that require time at varying scales between objects and agents. For instance, consider agent A and agent B on a soccer field. Suppose agent B leaves his favorite soccer ball on the field and goes on a bathroom break and agent A is left to practice with B’s soccer ball. However, we see Agent A use all of his effort to kick the soccer ball out of the field, disappearing from view. Agent A leaves followed by B returning to the field with a confused look. In this example, we have immediate causal physical interactions: Agent A exerts a lot of effort behind his kick, so we expect the ball to roll a far distance out of view. But there is also a concept of mentalization with Agent B’s confusion that occurs across a larger timescale. As discussed earlier, the specialization of these perception abilities and the temporal hierarchies at which they occur at fundamental aspects of cognition.

Second, no previous work (a) projects multimodal embeddings into LLM’s token decoder space for use in exposing model reasoning and (b) map respective hidden states of the decoder blocks onto human brain data. Since LLMs are autoregressive by nature and pretrained for causal generation (Daniel & Martin, 2024), probing how they can co-attend to visual tokens and system prompts (e.g. queries or dialogue about a given video) may offer secondary insights. Finally, there is a serious gap in research that attempts probing how vision models may separately encode dorsal and ventral voxel activations implicated in commonsense physics and social perception. Although the discussed evidence for this dissociation is clear in humans, current models are disproportionately focused on the ventral hierarchy (Ricci & Serre, 2020). This should not be interpreted as suggesting that explaining ventral topography is insignificant. Nonetheless, demonstrating that joint vision-language models can at least partially encode both streams during different task demands may be just as insightful. Indeed, “dual stream” ML models that separately learn “Where” pathways of spatial attention and “What” pathways of local features around eye fixation show robust evidence in separately encoding both hierarchies (Choi et al., 2023). If multimodal video-language models can uniquely encode dorsal voxels engaged by physical violations of expectation and ventral voxels engaged by psychological violations, then there is a clear upshot: it may hint that linguistically grounded video models are flexible representers of both (a) visual action and expected scene interactions and (b) the interaction of social agents with others and their environment.

## Methods

To pursue these questions, we create a novel spatiotemporal LLM training and encoding paradigm that projects video-level CLIP features into the token decoder block through a learnable linear layer. We use 3D video rendering scripts from both the ADEPT and AGENT datasets to create two separate training datasets of 10,000 videos that do or do not violate psychological or physical expectations. Spatiotemporal features are extracted by using a pretrained CLIP L/14 model that computes spatially and temporally pooled representations across all video frames (figure 1). Spatial representations are obtained by averaging all patch embeddings across each frame in the video to obtain a single vector representation. The resulting vectors of each frame are concatenated to achieve a video-level spatial representation. Temporal representations are acquired by averaging the frame-level embedding across the number of frames in the video.

**Figure 1.**
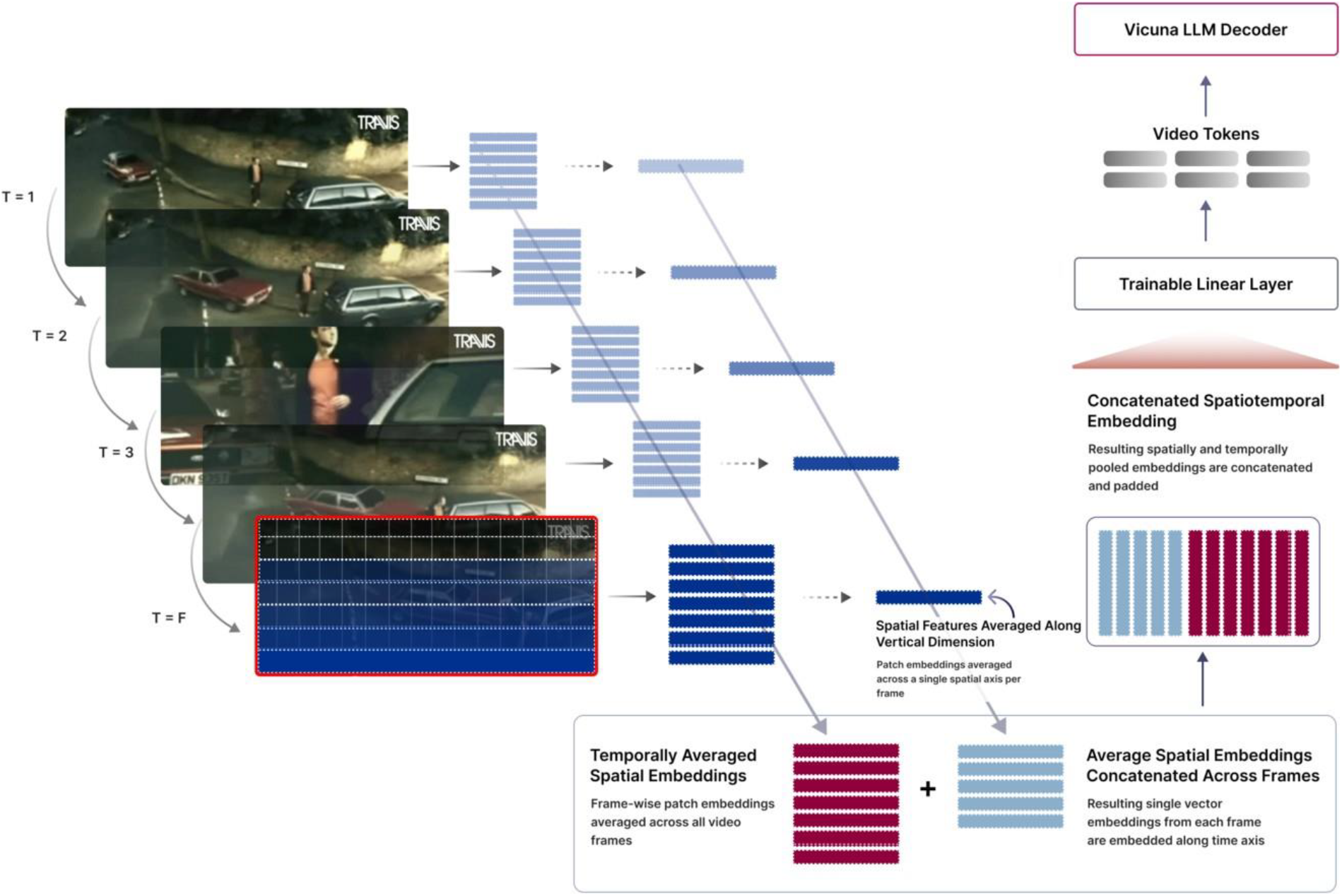
Original training paradigm using CLIP L/14 for spatiotemporal embedding extraction and LLM token projection CLIP L/14 (i.e. 14 patches) embeddings are first averaged along the y axis of the spatial dimension, retrieving a single spatial vector per video frame. These are then concatenated to form a video-level spatial representation. Temporal embeddings are retrieved by averaging the frame-wise spatial embeddings (unaveraged) along all frames.

Eventually, spatiotemporal embeddings are generated by concatenating padded spatially pooled representations with temporal averaged features. The training paradigm is based on video instruction tuning, using question-answer pairs for each video across psychological vs. physical expectations. For each training video, the system prompt (i.e. “question”) is concatenated with video-level tokens. The ground truth “answer” is a simple “yes” or “no.” These video-level tokens are computed through a simple linear projection matrix W that keeps both the visual encoder and LLM weights frozen while maximizing the likelihood of the target answer given both the visual token and system prompts. The original system prompt is *“Is this video surprising or not? You are to strictly reply with ‘yes’ or ‘no*.*’ “*” VID_TOKENS. This slowly projects the original spatiotemporal CLIP embeddings into a token space that can be exploited by the LLM in question answering.

Following two step training and finetuning, we create separate ridge regression models to predict voxel activity in a holdout dataset of one subject that undergoing fMRI scanning when viewing videos that violate psychological and physical expectations from the AGENT and ADEPT datasets, respectively (figure 2) (Liu et al., 2024). Broadly, the two encoding models are (1) Physical VoE which is fitted to only encode voxel activity for videos that do or do not violate physical expectation and (2) Psychological VoE which is only fitted for psychological expectation events. For each model, we obtain several features including (a) the *pre*-projected CLIP embeddings, (b) hidden states from layer 32 of the LLM’s decoder, and (c) corresponding logits. For each time point in a block of events that mapped to either a positive or negative psychological VoE, each of the aforementioned input features was separately fitted based on the VoE class it belonged to which derives from the given video sample that the multimodal LLM processes during inference. Encoding results are obtained using K-1 (K=10) fold validation across all BOLD timeseries vectors in a given sample. This results in a (50 x 70 x 70) x *num_timepoints* shaped matrix of explained variance, Pearson correlation, and mean absolute error (MAE) metrics. These scores are flattened across the temporal dimension (i.e. to obtain a shape of 50 x 70 x 70 voxel metrics) which results in a representative shape of metrics that can then be transformed into a cortical flat map.

**Figure 2.**
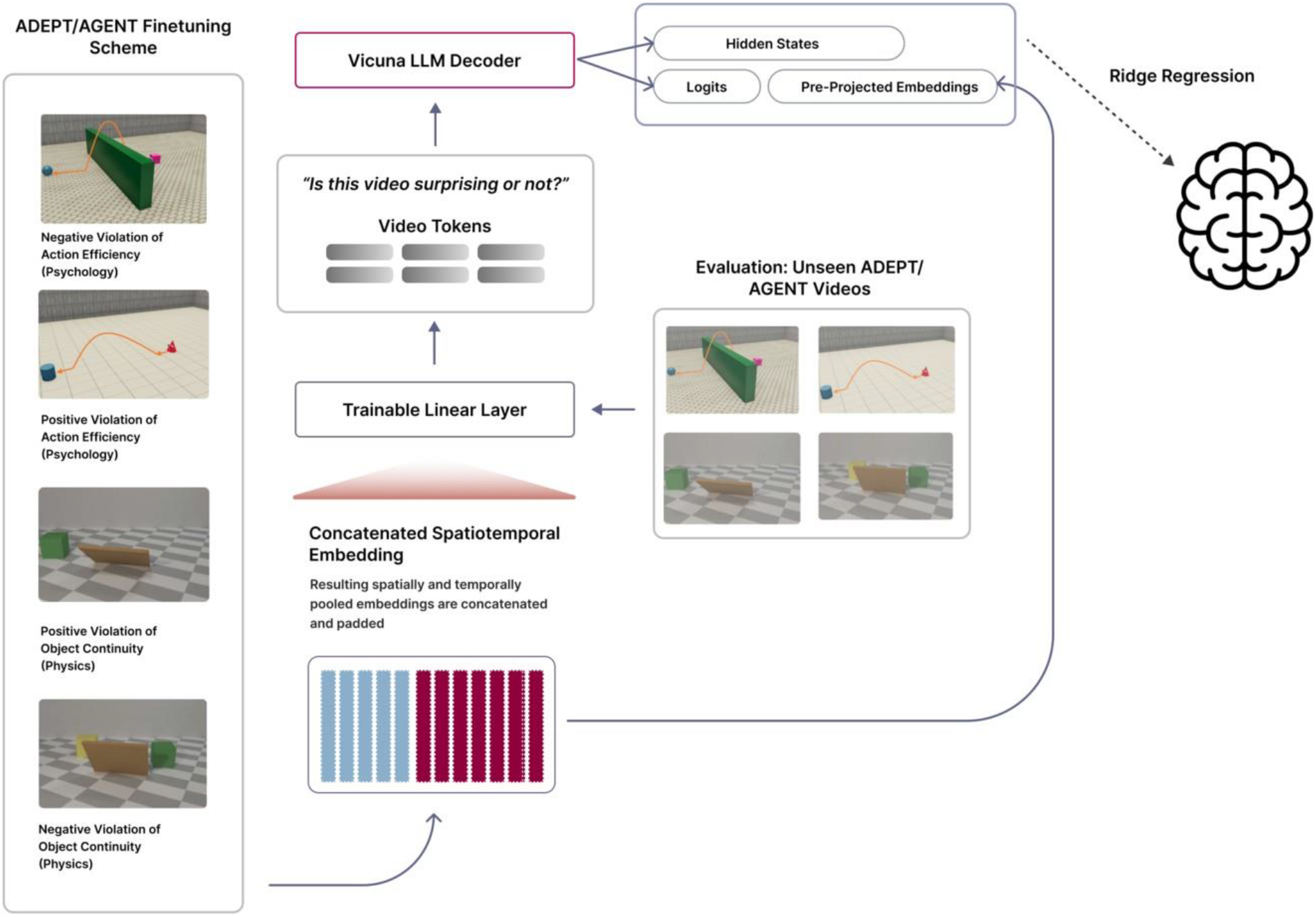
Modified AGENT/ADEPT dataset finetuning scheme. 10,000 video clips are generated containing approximately 1:1 split between samples that do/do not violate expectation in each of the psychology and physics tasks. Spatiotemporal features are extracted for each frame in all training videos while a trainable linear layer maximizes alignment with natural language tokens. The projection layer is frozen after training and used during evaluation on unseen ADEPT and AGENT videos to obtain visual tokens which are then projected into LLM decoder space. The LLM is prompted to answer whether the video is surprising or not based on the projected tokens. Finally, hidden states and logits are extracted from the decoder as it performs inference on each video. The original pre-projected embeddings are also fit into the regression model. All features are used to create separate voxel wise encoding models for a subject viewing the same videos.

## Results

### Multimodal LLMs Encode Physical and Psychological Violations of Expectations

We create two separate encoding models for psychological and physical VoE using only the hidden states and logits from the decoder block of the Vicuna 1.1 model during inference. These vectors are first flattened into a matrix that matches the spatiotemporal dimensions of the fMRI data and then used as predictors to fit the model in a K-1 (K=10) cross-validation scheme. When using hidden states alone, our encoding model can predict BOLD activity along sparsely distributed regions across the inferior-parietal lobule. Moreover, this model explains high variance in voxel wise activity (see figure 3. Redder is better) approaching R^2 values of 60%. Additionally, logits from the decoder are also obtained by multiplying the last hidden state matrix by decoder weights. After fitting prediction logits, we find that the regions of explained variance occur in effectively the same areas (Figure 4). However, on average, R^2 values rise by 10% (i.e. 70%). This effect of logits intensifying variance explained is also found when encoding for psychological VoE. Moreover, Pearson Correlation coefficients between predicted and actual voxel activities were comparably high in the same regions, consistent with R^2 scores (figure 5).

**Figure 3.**
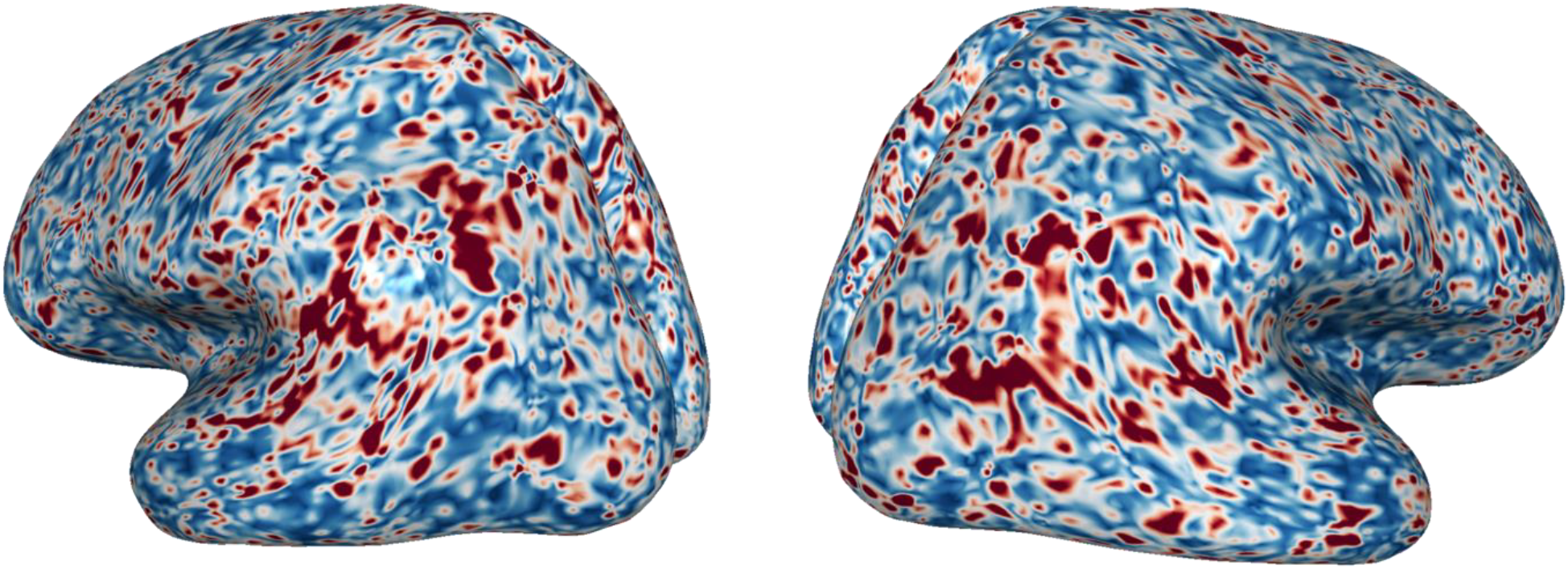
Explained variance (R^2) when **hidden states** encode for physical VoE only (0 - 0.6)

**Figure 4.**
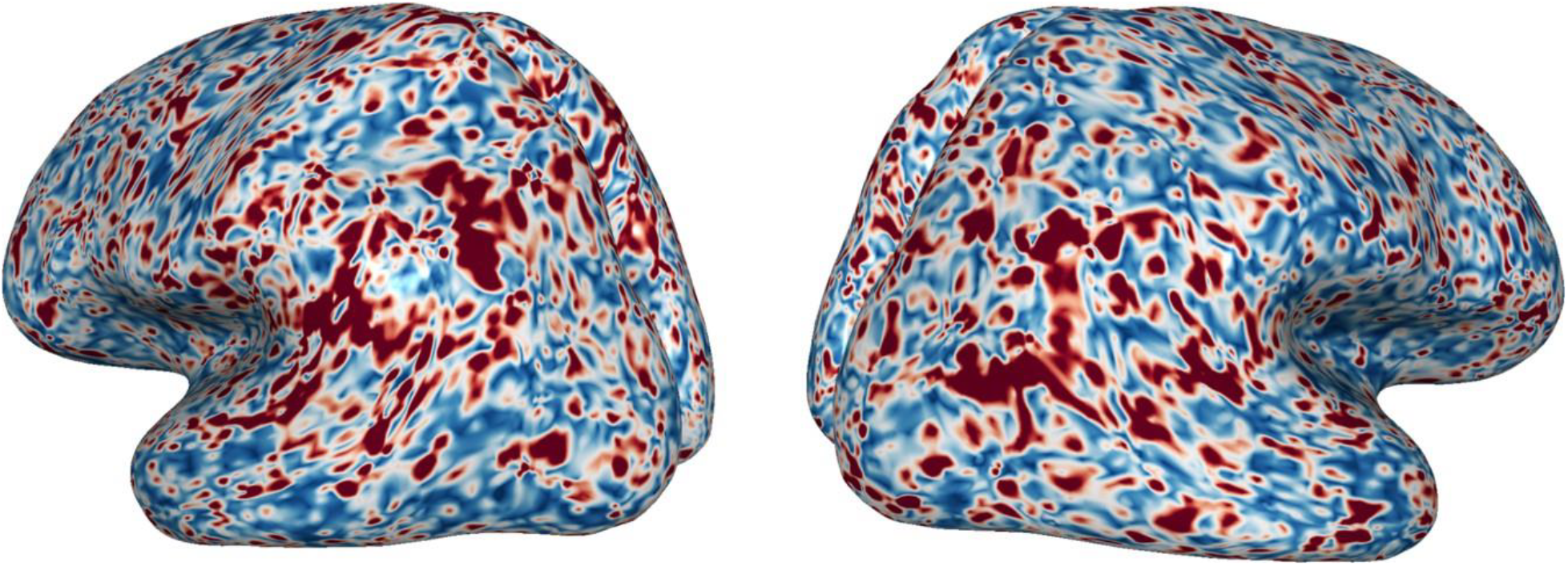
Explained variance (R^2) when **logits** encode for physical VoE only (0 - 0.7)

**Figure 5.**
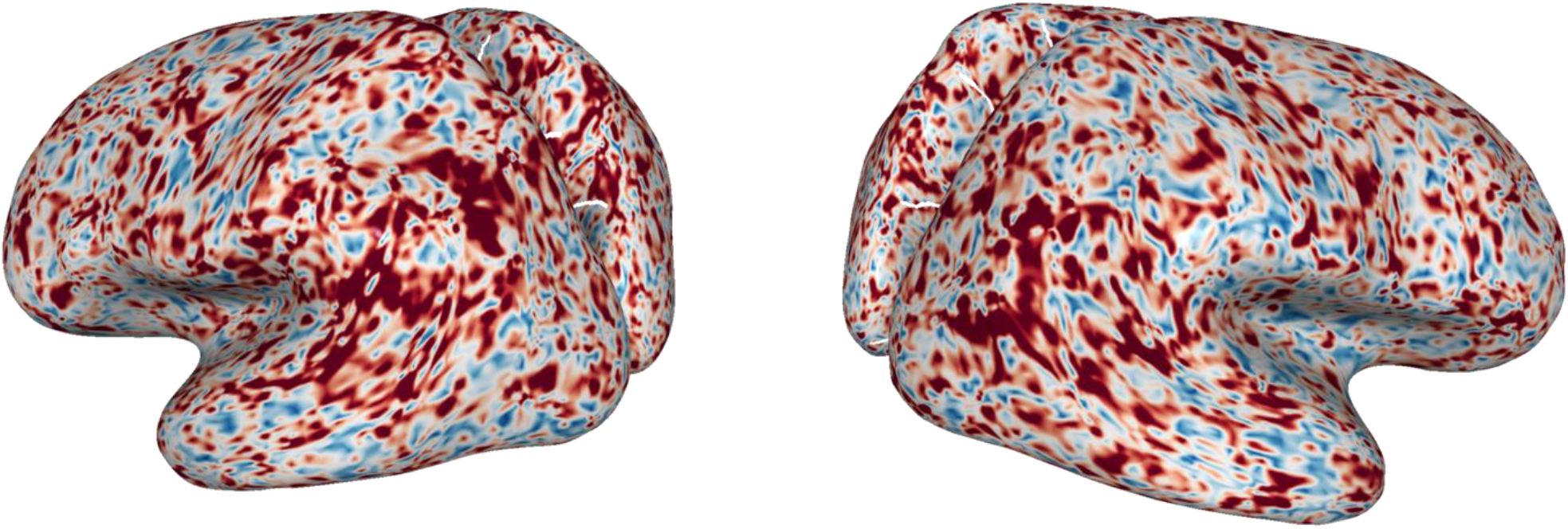
Correlations between predicted and actual voxel activations when **logits** encode for physical VoE only (0 - 0.7)

We perform the same encoding paradigm for psychological violations of expectation. Surprisingly, hidden states and logits encode *highly* distributed regions of voxel activity extending into the lateral visual pathway and more extended parietal areas (Figure 6). Notably, although these hidden states encode overlapping dorsal regions, they also predict ventral areas that cannot be explained or predicted as adequately in the physical VoE model (Figure 7).

**Figure 6.**
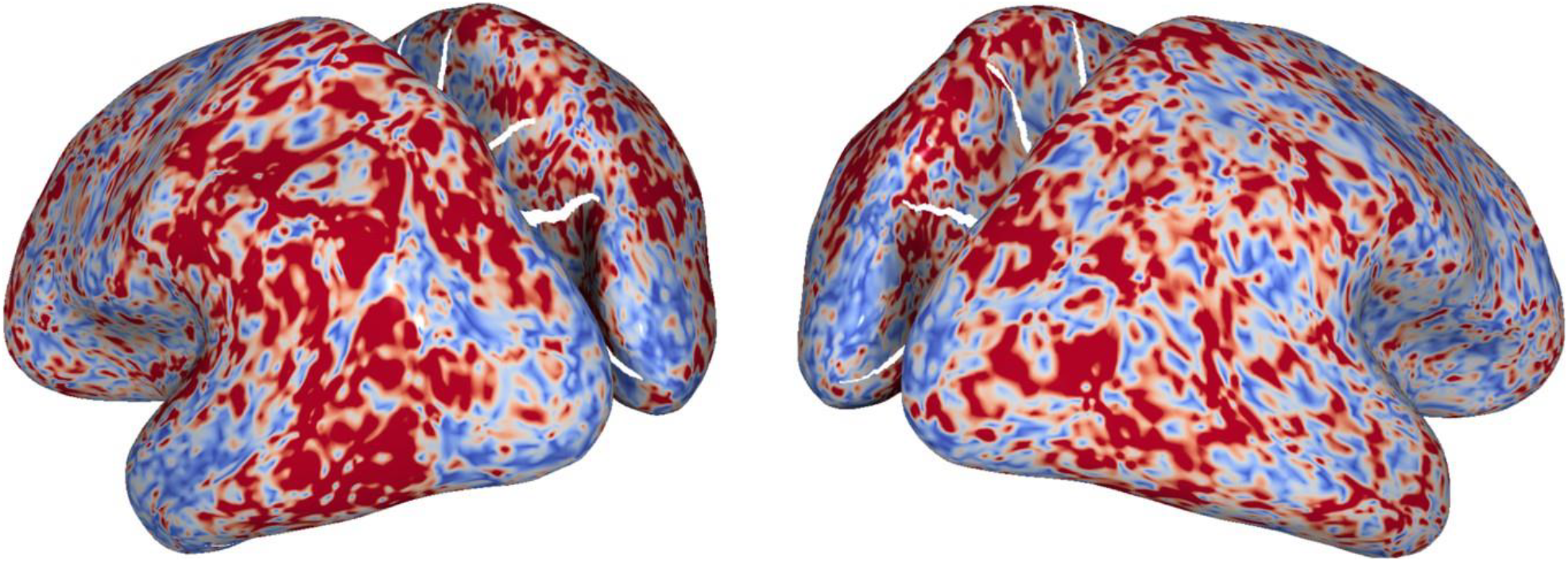
Explained variance (R^2) when **hidden states** encode for psychological VoE only (0 - 0.6)

**Figure 7.**
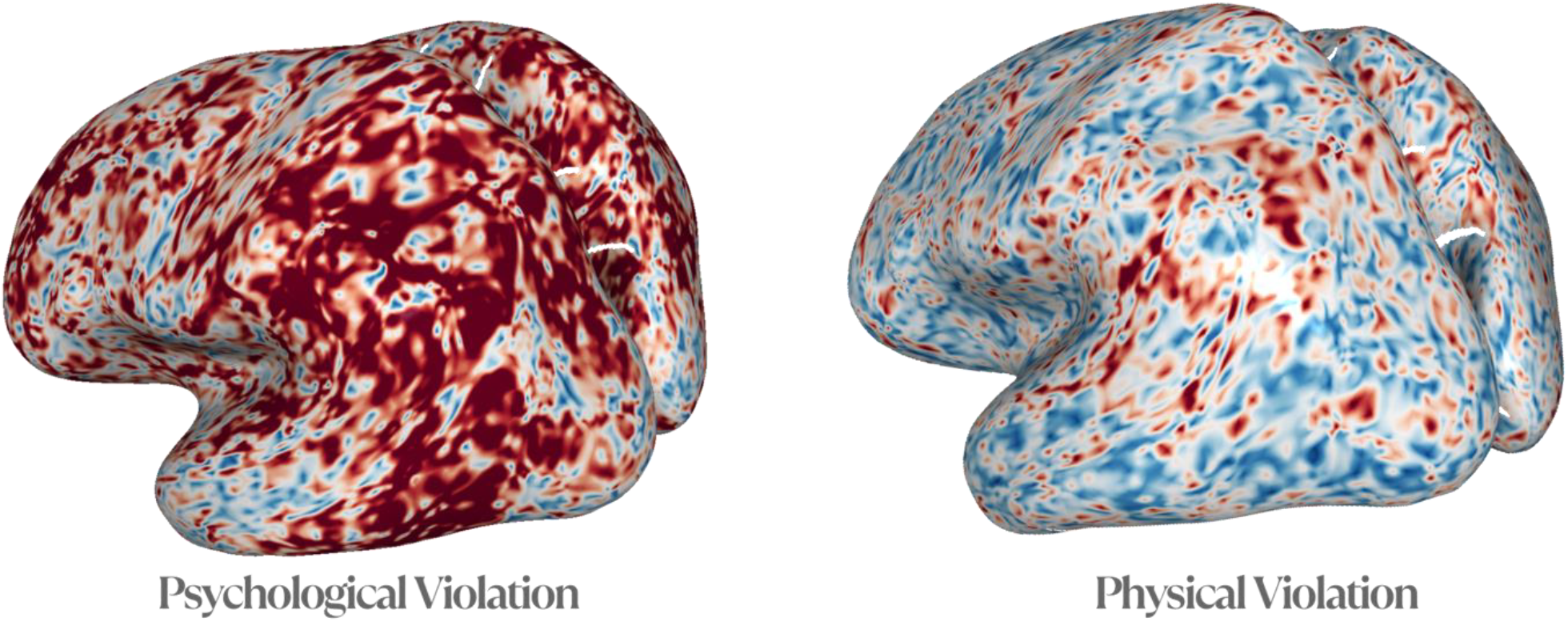
Comparison of Pearson correlations for psychological VoE and physical VoE encoding models using **hidden states** (0-0.6)

### Pre-Projected Video Embeddings Explain Similar Variance with Hidden States for Physical but not Psychological

As noted in figure 2, pre-projected spatiotemporal CLIP embeddings represent pooled multimodal representations that have not yet been projected into the LLM token’s decoder space. It was a natural next step to explore how voxel encoding performance may or may not change after these embeddings are aligned with text token representations in the decoder of the LLM’s last layer. The motivation for this comparison was simple: although CLIP embeddings are inherently vision-text aligned, it is plausible that – in their pre-projected form – they represent a simple concatenation of frame-level patches that are not yet accessible to for semantic reasoning. Thus, we hypothesized that the distribution of explained variance in voxel activity may benefit from LLM projection for psychological expectations more than physical expectations. This is grounded in the premises that (1) temporal frame embeddings may still represent artifacts of motion and object/scene changes and (2) the intuitive physics network may not access language representations as heavily as social perception. Indeed, after creating separate encoding model schemes for pre-projected embeddings and hidden state representations, we find two core results. First, token projection does not benefit voxel predictivity for intuitive physics but does benefit predictivity for intuitive psychology. In other words, the pre-projected video embeddings explain nearly identical magnitude and regions of variance for intuitive physics but not psychology. Second, for psychological expectations, hidden states encode a large distribution of voxel activity that extend into upstream dorsal and ventral regions, including V3, V4, IT, and STS (figure 8).

**Figure 8.**
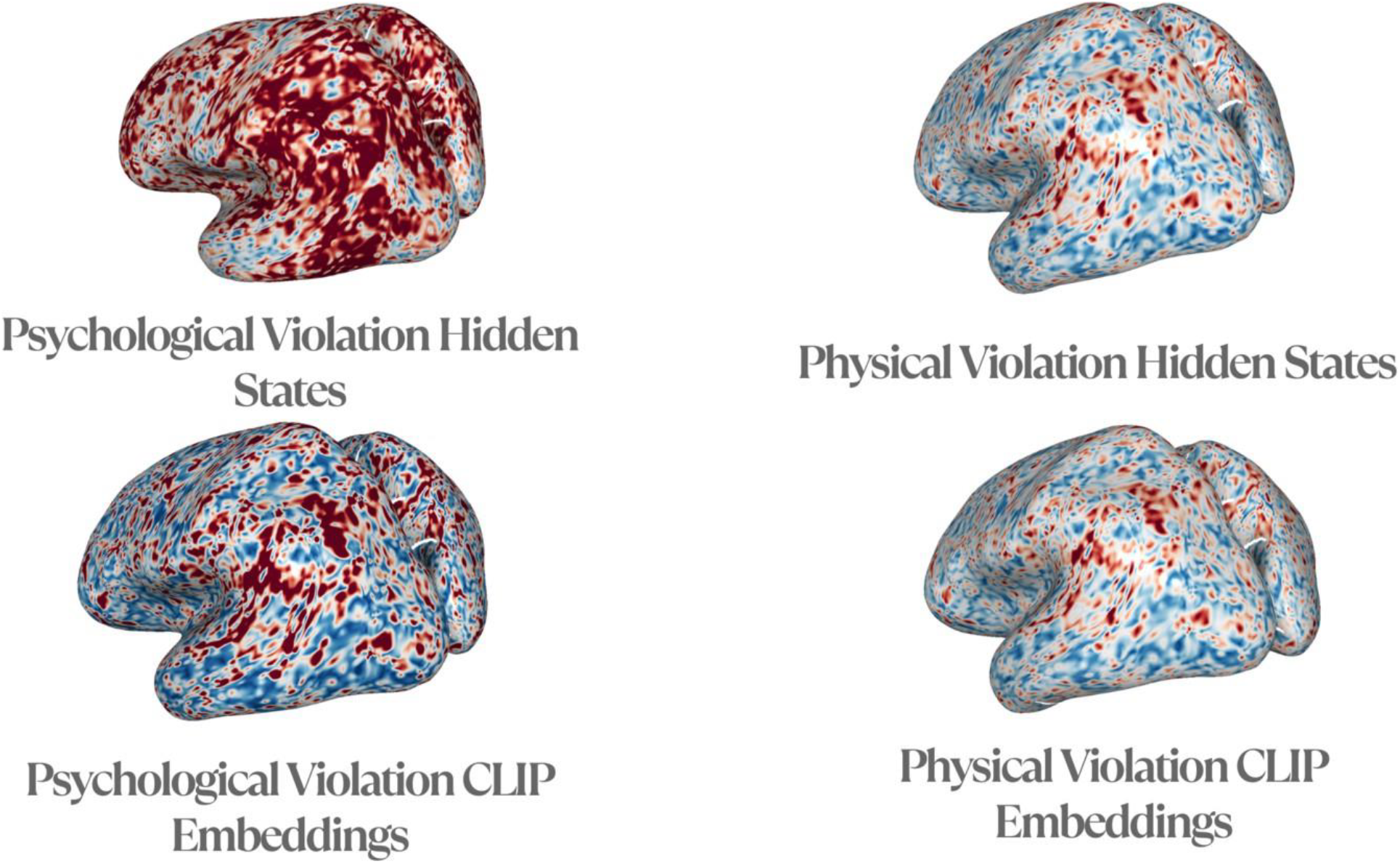
Explained variance (R^2) score distributions (0-0.6) between pre-projected CLIP embeddings and hidden states in psychology vs. physics encoding models. Left column demonstrates change in explained variance only when encoding psychological violations between hidden states and CLIP embeddings. Right column demonstrates similar areas and magnitude of explained variance for physical violations.

## Discussion

Navigating and perceiving a world with an ever-changing gradient of visual concepts, social interactions, and physical events is an exceptional feature of human cognition. The brain’s ability to flexibly reason in a multimodal manner while still being hierarchically sensitive to the timescales at which these events take place is a property that many modern deep learning models scrape by in answering. More importantly, the dual stream topography and dissociation underlying our intuitions about the physics and psychology of agents has been negligibly answered by the current crop of AI models. In this paper, we deployed a novel vision-language model that learns to project spatiotemporal vision embeddings into the decoder’s token space, aligning video-level representations into a format that can be exploited in an autoregressive manner for downstream reasoning. We finetune this video-text token alignment by projecting video and question-answer language pairs when viewing psychological and physical violations in both the AGENT and ADEPT datasets, respectively. Our voxel wise encoding paradigm marks a first attempt at aligning both pre-projected visiolinguistic embeddings and hidden state representations with brain activity when viewing violations of psychological and physical expectation. Two core results emerged from this analysis: First, we find that in the same model (i.e. with the same, frozen linear projection layer), hidden state and logits representations from the last decoder block begin to explain distinct regions of variance divided between (a) highly distributed activations in ventral and ventral-dorsal pathways when encoding for social actions and (b) minimal, sparse activations in inferior-parietal dorsal areas. Second, this divergence in ventral psychological vs. dorsal physical expectation encoding only clearly manifests after CLIP embeddings are projected into decoder tokens. In conjunction, only the psychological encoding model benefits from video token projections in terms of explaining more distributed ventral-dorsal activations while the physics model does not.

Why might hidden state activations more reliably benefit voxel predictions for psychological VoE but not physical? We propose that this distinction arises by virtue of the different functional necessity for language-aligned visual understanding in both tasks. The lateral visual pathway, which is theorized to integrate low-level features in the early visual cortex (EVC) and high-level features in the STS, is increasingly multimodal in its association. Previous research found that the spatial topography of neurons in the lower bank of the STS finds cells that respond to more than one modality (Zheng et al., 2010). Past experiments also more generally suggest that multimodal representations in the ventral visual pathway are present. For instance, single neuron recordings of the medial temporal lobe (MTL) show increasing multimodal invariance to representations of the same individual presented across different sensory modalities (Quiroga et al., 2009). In contrast, the dorsal cortex may not need be penetrated by top-down semantic concepts in the same manner that feedback projections in the ventral stream serve. It is theorized that dorsal pathways may compute spatial relationships among object parts without needing to access the form and linguistically grounded concepts of the parts themselves. This information is then propagated to the ventral pathway to support object recognition (Ayzenberg & Behrmann, 2022). This may explain why we find highly distributed ventral-dorsal regions of explained variance when using hidden state representations in psychological expectations. Although stimuli with psychological violations contained agentic interactions, they are still physically grounded in the sense that spatial relations are still being computed to build up relationships between agents and the environment.

Finally, what are the implications of aligning multimodal AI models with separable processing streams of the brain? As noted, human perception is exceptional in the sense that its topography has naturally evolved to serve processing at varying timescales of integration and inference with energy efficient receptive field structures (Vincent et al., 2005). Although the biological plausibility of these networks is not within the scope of this paper, future research ought to explore the temporal receptive windows (TRWs) of modern video-language transformers and probe the receptive field structures of neurons across layers of vision-language models. Creating AI systems that can accurately and rapidly guide their actions while jointly understanding the goals of other human and non-human agents widens the possibility for the development of empathetic systems. For instance, imagine the following scenario: a socially intelligent robot operates a rollercoaster. They see a child hastily running to their seat in an overexcited manner. This robot may understand that this excess goal leads to the child overlooking safety harnessing. Physical intuitions about the motion of their safety harness can confirm these social evaluations and allow the robot to plan its safety protocols accordingly. Finally, it should be noted that this paper uses simple video stimuli while CLIP was originally trained on a mix of four hundred million image-text pairs in the wild sourced from the Internet (Xu et al., 2023). Although this presents a limitation, current intuitive psychology and physics fMRI data has only been analyzed using computationally rendered samples as seen in the AGENT and ADEPT datasets. In this direction, future research should focus on conducting fMRI analyses with naturalistic stimuli that is coded with physical and psychological expectation annotations.

## Notes

### Competing Interest Statement

The authors have declared no competing interest.

